# Unraveling the Molecular Landscape of Uterine Fibroids, Insights into *HMGA2* and Stem Cell Involvement

**DOI:** 10.1101/2024.04.26.591351

**Authors:** Emmanuel N. Paul, Tyler J. Carpenter, Laura A. Pavliscak, Abigail Z. Bennett, Maria Ariadna Ochoa-Bernal, Asgerally T. Fazleabas, Jose M. Teixeira

**Affiliations:** Department of Obstetrics, Gynecology and Reproductive Biology, College of Human Medicine, Michigan State University, Grand Rapids, MI 49503, USA

**Keywords:** Myometrium, fibroids, stem cell, transcriptome, extracellular matrix

## Abstract

Uterine fibroids are prevalent benign tumors in women that exhibit considerable heterogeneity in clinical presentation and molecular characteristics, necessitating a deeper understanding of their etiology and pathogenesis. *HMGA2* overexpression has been associated with fibroid development, yet its precise role remains elusive. Mutations in fibroids are mutually exclusive and largely clonal, suggesting that tumors originate from a single mutant cell. We explored a possible role for *HMGA2* overexpression in differentiated myometrial cells, hypothesizing its potential to induce a stem cell-like or dedifferentiating phenotype and drive fibroid development. Myometrial cells were immortalized and transduced with an *HMGA2* lentivirus to produce HMGA2hi cells. *In vitro* stem cell assays were conducted and RNA from HMGA2hi and control cells and fibroid-free myometrial and HMGA2 fibroid (HMGA2F) tissues were submitted for RNA-sequencing. HMGA2hi cells have enhanced self-renewal capacity, decreased proliferation, and have a greater ability to differentiate into other mesenchymal cell types. HMGA2hi cells exhibit a stem cell-like signature and share transcriptomic similarities with HMGA2F. Moreover, dysregulated extracellular matrix pathways are observed in both HMGA2hi cells and HMGA2F. Our findings suggest that HMGA2 overexpression drives myometrial cells to dedifferentiate into a more plastic phenotype and underscore a pivotal role for HMGA2 in fibroid pathogenesis.

## Introduction

Uterine fibroids, or leiomyomas, represent the most prevalent benign tumors in women of reproductive age, affecting approximately 70-80% of women during their lifetime (1). These tumors are mainly characterized by extensive extracellular matrix (ECM) deposition and remodeling (2). Despite their high incidence, the etiology and pathogenesis of uterine fibroids remain poorly understood.

Uterine fibroids exhibit considerable heterogeneity in terms of their clinical presentation, growth patterns, and response to therapeutic interventions (3), which suggests different phenotypic subtypes. Recent reports showing molecular characterization of fibroids have provided unprecedented insights into the complex interplay of genetic, epigenetic, and cellular factors contributing to the development and progression of uterine fibroids (4–6) and support the classification of uterine fibroids into distinct subtypes based on their combined molecular and histological characteristics. Mutations identified in various genes, including *MED12*, *FH*, and *COL4A5*, have been associated with distinct subtypes of fibroids, potentially contributing to the diverse clinical manifestations observed in affected individuals (7,8). Unraveling the molecular mechanisms driving the pathogenesis of the various fibroid subtypes is critical for developing targeted therapeutic strategies and personalized treatment approaches.

Another genetic factor associated with fibroid development is overexpression of the high-mobility group AT-hook 2 (*HMGA2*) gene (9–12). *HMGA2* is located on chromosome 12q15 and encodes a non-histone chromatin-binding protein that plays a crucial role in regulating gene expression especially during embryonic development, including stem cell pluripotency, lineage specification, and organogenesis (13). *HMGA2* transcripts are detected in most embryonic tissues and progressively become absent in adult tissues, except for the testis, the lung, and the kidney (14). Its dynamic expression and regulatory functions are tightly orchestrated to ensure proper development, with aberrations in *HMGA2* expression linked to developmental abnormalities and diseases (13,15,16). Overexpression of *HMGA2* commonly resulting from the translocation between chromosome bands 12q15 and 14q24 has been previously described as the second major subtype of uterine fibroids (HMGA2F), constituting approximately 7% of fibroid cases (17,18).

The discovery of a potential stem/progenitor cell population within the myometrium has generated interest in uncovering the involvement of these cells in the initiation and growth of fibroids (19,20). Indeed, uterine fibroids are thought to be a clonal disease (21,22), and since most clonal diseases have a single cell origin, the prevailing hypothesis has been that a dysregulated myometrium stem cell could be the cell of origin for uterine fibroids (23). However, the molecular mechanisms governing the interplay between myometrium stem cells and *HMGA2* in uterine fibroids remain largely unexplored. In this study, we explored an alternative hypothesis regarding the cell origin for uterine fibroids by postulating that a mutated differentiated myometrial cell could give rise to a fibroid. We show that *HMGA2* overexpression in myometrium cells induces stem/progenitor cell properties, suggesting a potential gain of function for the gene in fibroid development. Such insights may pave the way for the development of novel therapeutic targets that can effectively address uterine fibroids in a subtype-specific manner.

## Methods

### Myometrium cell line generation

HEK293T cells were transfected with hTERT plasmid, Addgene #85140 (24) and helper plasmids (25) to prepare hTERT lentivirus. Myometrial cells were dissociated and isolated from a fibroid-free myometrium patient (MyoN) as previously described (26) and incubated at 37°C overnight in growth medium (DMEM/F12, 10% FBS) prior to transduction with lentiviral preparations. The subsequent day, transduced MyoN cells were selected with 50 µg/mL of hygromycin for 1 week. Following selection, MyoN-TERT cells were maintained in medium with 5 µg/mL hygromycin.

### *HMGA2* Overexpressing and Control MyoN-TERT Cell Lines

MyoN-TERT cells were suspended in 1% bovine serum albumin blocking buffer, incubated with a rabbit anti-human CRIP1 primary antibody (ThermoFisher, #PA5-24643). Cells were then incubated with an Alexa-647 anti-rabbit secondary antibody and washed with flow buffer, and resuspended in flow buffer with of 4′,6-diamidino-2-phenylindole (DAPI). Cells were sorted by the flow cytometry core at Van Andel Research Institute (VARI) using a FACSymphony S6 cytometer (BD Biosciences) and analyzed with FlowJo Software (BD Biosciences, version 10.8.1). Lentiviral particles isolated from in HEK293T cells transfected with either a custom *HMGA2* (VectorBuilder) or empty control plasmid (VectorBuilder) were used to transduce CRIP1^-^ MyoN-TERT cells as described above. Selection of successfully transduced MyoN-TERT live cells expressing GFP were sorted by FACS. *HMGA2* overexpressing cells are called HMGA2hi cells.

### Proliferation Assay

HMGA2hi and control cells were plated in triplicate a 96 well plate at 2,000 cells/well, and cell proliferation was recorded every 4 h using an Incucyte SX5 instrument (Sartorius), following the manufacturer’s protocols, until cell confluence was reached. Cell proliferation was quantified using GFP and phase contrast images at each time point with the cell-by-cell software module of the instrument. The experiment was repeated three times using three different cell passages.

### Colony Formation Assay

Matched passages of HMGA2hi and control cells were plated at 50 cells/cm^2^ in triplicate overnight, then switched to MesenPro RS (Thermo Fisher, # 12746012) for 2 to 3 weeks. Cells were cultured in different wells of the same plate, and both cell types were assayed on the same day. Cultures were fixed in 4% paraformaldehyde (PFA) and stained with crystal violet for colony visualization. Images were taken using a Nikon SMZ18 microscope and Ds-Ri1 camera (Nikon Instruments Inc.). Colonies were counted and total surface area was estimated using ImageJ (version 1.53k), and the percent colony-forming units (CFUs) was calculated as (number of colonies/number of cells plated) × 100 and averaged for triplicates.

### Differentiation Assays

For osteogenic and adipogenic differentiation, matched passage HMGA2hi and control cells were plated in duplicate at 50% to 80% confluency in growth media overnight and then cultured for 10 days in fresh StemPro Adipogenesis Differentiation (Thermo Fisher Scientific, #A1007001) or StemPro Osteogenesis Differentiation (Thermo Fisher Scientific, #A1007201) media according to the manufacturer’s instructions. Cells were cultured in regular growth media to serve as differentiation controls. To assay adipogenic differentiation, cultures were fixed in 4% PFA and stained using Oil Red O (Sigma, #01391) according to the manufacturer’s instructions. To assay osteogenic differentiation, cultures were stained for alkaline phosphatase activity using the Alkaline Phosphatase (AP), Leukocyte kit (Sigma, #86R-1KT) according to the manufacturer’s instructions.

### 3D Culture and Trichrome Staining

60,000 cells of HMGA2hi and control cells, as well as primary MyoN and fibroid cells, were seeded in scaffold-free 96-well agarose micro-molds (MilliporeSigma, Z764043), following the protocol outlined by Wiwatpanit et al. (27). Spheroids were cultured for 4 days in growth medium. Following culture, agarose micro-molds were sealed with agarose and fixed in 4% PFA at 4°C overnight. Subsequently, they were briefly rinsed in PBS and stored in 70% ethanol at 4°C until processed for standard paraffin embedding and sectioning at 6 μm. Spheroid sections were stained with trichrome stain according to the manufacturer’s protocol (StatLab #KTTRBPT). Briefly, sections were deparaffinized with two xylene baths and rehydrated with three absolute ethanol washes. After hydration, spheroid sections were incubated in Bouin’s fluid at room temperature (RT) overnight. The next day, after rinsing in running tap water, slides were stained with Mayer’s hematoxylin and rinsed in running tap water. Spheroid sections were then immersed in the one-step trichrome stain and quickly rinsed in running tap water. Slides were dehydrated through three changes of absolute ethanol, cleared in three changes of xylenes, and a coverslip was applied using Permount mounting medium (Fisher Scientific #SP15). Collagen deposition was determined using the color deconvolution ImageJ plugin (version 1.53k). Relative collagen area was calculated by the ratio of blue stain (collagen) over the red stain (cytoplasm).

### Tissue Collection and *HMGA2* fibroid mutation detection

The use of human tissue specimens was approved by the Spectrum Health Systems and Michigan State University Institutional Review Boards (MSU IRB Study ID: STUDY00003101, SR IRB #2017-198) as secondary use of biobank materials. Fibroids samples (HMGA2F, n = 23) and fibroid-free myometrial (MyoN, n = 32) from pre- menopausal women (aged 28-55) were obtained following total hysterectomy. All patients who participated in the study gave consent to donate tissue through the Spectrum Health Biorepository. Human samples were processed as previously described (4). Briefly, samples were washed with phosphate-buffered saline, minced, and immersed in RNAlater (Sigma, Saint Louis, MO, USA) for RNA-seq analyses. *HMGA2* fibroid (HMGA2F) expression was determined by RT-qPCR using the iScript Advanced cDNA Synthesis Kit (BioRad, #1725037) for the reverse transcription and *HMGA2* PrimePCR primer (BioRad, #qHsaCED0021425) for the qPCR. RPL17 was use as a reference gene with forward ACGAAAAGCCACGAAGTATCTG and reverse GACCTTGTGTCCAGCCCCAT primers.

### RNA Isolation, Library Preparation and Sequencing

Total RNA from HMGA2hi and control cells was isolated from 3 different passages using an RNeasy mini kit (Qiagen) and stored at −80 °C in nuclease-free water. RNA integrity values from cells and tissues were determined with an Agilent 2100 Bioanalyzer (ThermoFisher), and values ≥7.5 were used for library preparation and paired-end (2 × 100 bp) RNA-sequencing on an Illumina NextSeq 6000 instrument (Illumina). Libraries were prepared using a Kapa RNA HyperPrep kit with ribosomal reduction, pooled, and sequenced on flowcells to yield approximately 50–60 million reads/sample. Raw fastq files were deposited in the NCBI Gene Expression Omnibus (GSEXXXX).

### RNA Seq Analysis

RNA-seq reads were trimmed for quality and adapters using TrimGalore (version 0.6.4), and quality trimmed reads were assessed with FastQC (version 0.11.7). Trimmed reads were mapped to Homo sapiens genome assembly GRCh38 (hg38) using STAR (version 2.6.0c). Reads overlapping Ensembl annotations (version 99) were quantified with STAR prior to model-based differential expression analysis using the edgeR-robust method. Genes with low counts per million (CPM) were removed using the filterByExpr function from edgeR (28). Scatterplots of two selected principal components were constructed with the PCAtools R package (version 2.12.0) to verify sample separation prior to statistical testing. Generalized linear models were used to determine if principal components were significantly associated with cell type. Differential expression was calculated by comparing HMGA2hi to control cells and HMGA2F to MyoN tissues and differentially expressed genes (DEGs) were visualized with a volcano plot generated using the EnhancedVolcano (version 1.18.0) package in R. Boxplots of the log_2_(CPM) values were generated using the R package ggplot2 (version 3.4.4). Venn diagrams of the overlapping DEG of the cells (HMGA2hi vs. control) and DEG of tissues (HMGA2F vs. MyoN) were constructed using the eulerr package (version 7.0.0). Gene set enrichment and overrepresentation analyses were completed with clusterProfiler package (version 4.8.2) using MSigDB (29) and DOSE (30) reference databases.

### Statistics and Reproducibility

Bioinformatic statistics were performed using the listed packages in R (version 4.3.2). DEGs in the RNA-seq of HMGA2hi vs. control and the HMGA2F vs. MyoN were identified as those having a Benjamini–Hochberg FDR corrected p <0.05 (31). Data with unequal variances were log-transformed, and the homogeneity of variances was verified before the completion of analyses. Hypergeometric testing was performed using the function phyper from R. For the colony formation assays, comparison of two means was performed with a two-sided student t test, and significance was determined at p <0.05 after confirming normal distribution using GraphPad Prism (version 10.2.2).

## Results

### *HMGA2* overexpression increases myometrium cell plasticity

Our initial objective was to examine whether *HMGA2* overexpression enhances stem cell/progenitor-like properties in a differentiated myometrium cell line. Primary myometrial cells were immortalized (MyoN-TERT) and subsequently depleted for stem/progenitor cells via fluorescence-activated cell sorting (FACS) (Figure 1a and 1b), utilizing the cell surface marker CRIP1, a putative myometrium stem/progenitor cell marker recently identified in our laboratory (26). Among the MyoN-TERT cells, 73.7% of the MyoN-TERT were CRIP1^-^ (Figure 1b) and were sorted for downstream experiments. CRIP1-depleted MyoN-TERT cells were then transduced with either *HMGA2* (HMGA2hi) or a control virus. We performed proliferation, colony formation and differentiation assays to determine stem cell-like properties induced by *HMGA2* overexpression. The proliferation assay revealed reduced cell proliferation in HMGA2hi cells compared to the control cells (Figure 1c). Quantitative analysis of the colony formation assay showed a significant 4.3-fold increase in the number of cells with self-renewal capacity in the HMGA2hi cells (mean = 128) compared to control cells (mean = 30) (Figure 1d). While both control and HMGA2hi cells exhibited lipid droplets when cultured in adipogenic media (Figure 1e), indicating their ability to differentiate into adipocytes, staining in the osteogenic differentiation assay revealed alkaline phosphatase activity only in HMGA2hi cells and not in control cells when cultured in osteogenic media (Figure 1f).

**Figure 1.**
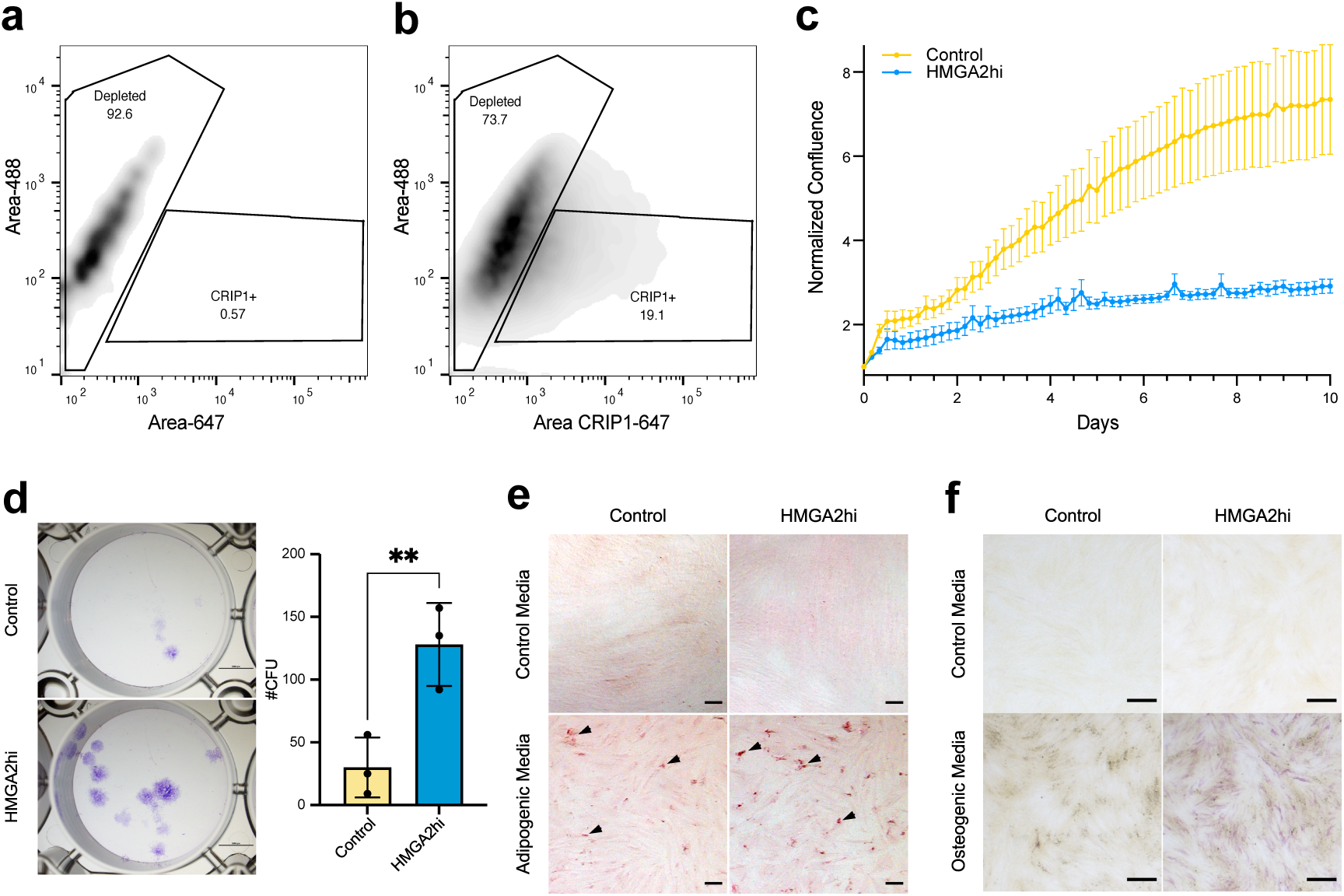
*HMGA2* overexpression increases myometrial cell plasticity. Scatter plot of the gating strategy used for cell sorting of depleted stem/progenitor MyoN-TERT, using the unstained sample as control **(a)** and the full stain sample to isolate the depleted cell population **(b)**. Quantification of the cell proliferation from the cell-by-cell function of the Incucyte, measuring the area occupied by cells (cm^2^/image) for each image scan normalized to time = 0 (n = 3, error bars represent standard of the mean, sem) (**c**). The yellow curve represents the control MyoN-TERT cells, and the blue curve the HMGA2hi cells, error bars represent standard error of the mean. Representative images of colonies formed by the control and HMGA2hi cells and plot of colony forming efficiency represented as %CFUs (#CFU/cells seeded × 100) (n = 3, error bars = sem) **(d)**. Representative images of control and HMGA2hi cells grown in control growth media and adipogenic **(e)** or osteogenic **(f)** differentiation media (n = 3). Adipogenic and control cultures were stained with Oil Red O (red color, black arrows), and osteogenic and control cultures were stained for alkaline phosphatase activity (purple color). Scale bar for the adipogenic and osteogenic assays are 500 µm. **p<0.01; by two-tailed student t-test.

### Stem cell-like transcriptomic signature is induced by *HMGA2* overexpression in myometrial cells

Principal component analysis (PCA) plot of HMGA2hi and control cells revealed sample separation primarily along principal component (PC) 2, which accounted for 29% of the variance, indicating that *HMGA2* overexpression induces significant transcriptomic changes in the immortalized myometrial cells (Figure 2a). Notably, the most prominent separation observed in PC1, representing 49% of the variance, was associated with cell passage and was thus incorporated as a co-variable for subsequent analyses. A volcano plot depicted a total of 709 DEGs with an FDR < 0.05 in HMGA2hi compared to control cells (Figure 2b). Among these, 304 genes were downregulated, while 405, including *HMGA2* (Log_2_FC = 2.2, FDR = 3.4 x 10^-55^), were upregulated (Table S1). Additionally, *CSPG4*, a gene associated with mesenchymal stem cells (26), was significantly upregulated (Log_2_FC = 0.4, FDR = 3.1 x 10^-2^) in HMGA2hi cells (Figure 2c). However, other putative myometrial stem/progenitor-related genes such as *PDGFRβ* (32), *SUSD2* (32), *MCAM* (32), *CD44* (19), and *CD34* (33) did not exhibit significantly (FDR > 0.05) higher expression in the HMGA2hi cells compared to control cells, except for *ITGA6* (also known as *CD49f*) (33), which was downregulated (Figure S1). Gene set enrichment analysis using the MSigDB (29) for cell type signature gene sets (C8) identified 62 significant pathways (FDR <0.05) (Table S2), including two upregulated stem cell gene sets (Figure 2d).

**Figure 2.**
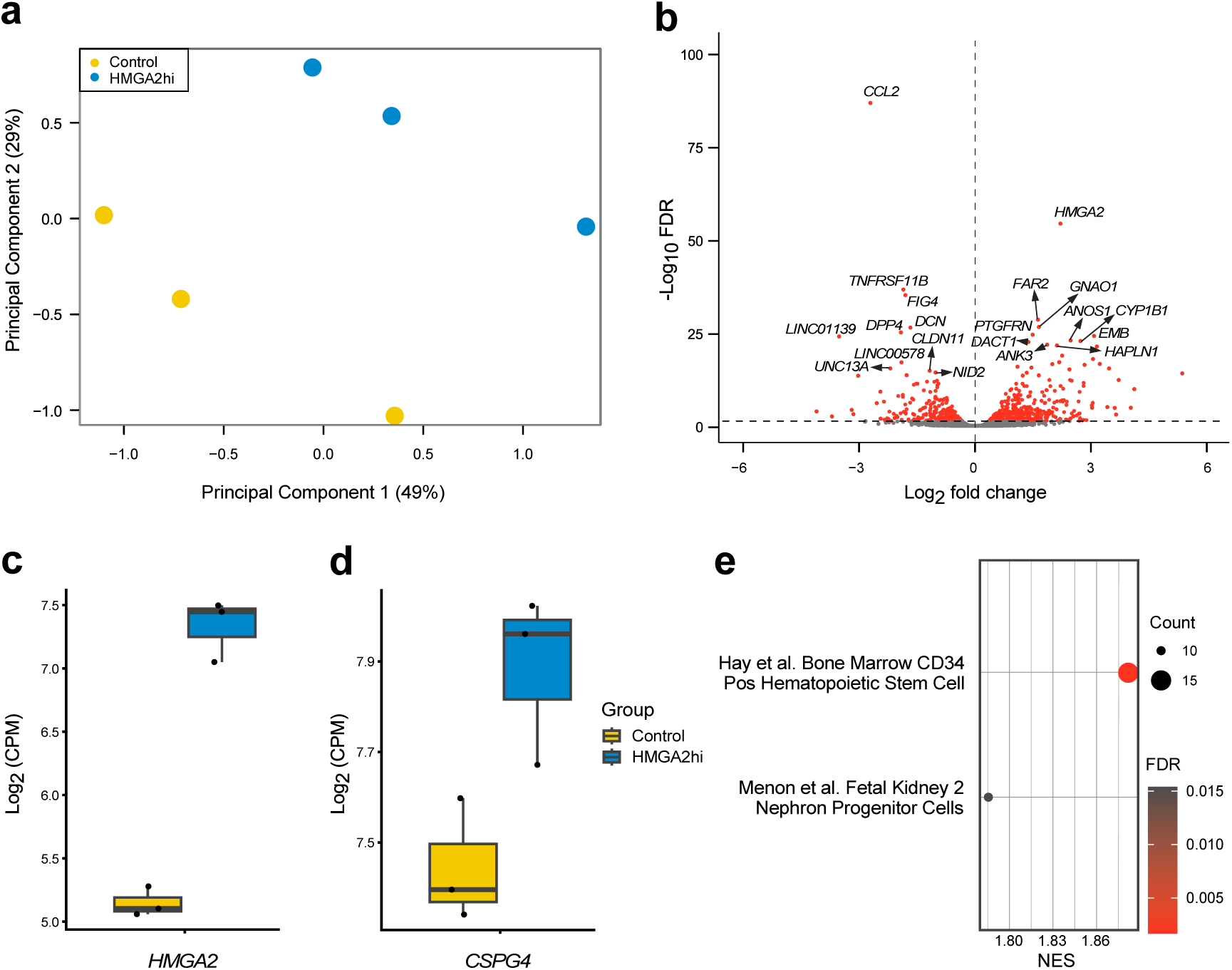
HMGA2hi transcriptome is enriched for stem cell markers and pathways. Principal component (PC) plot analyses for PC1 vs PC2 shows samples for control and HMGA2hi cells (n = 3/group) in two-dimensional space. Each dot represents one sample **(a)**. Volcano plots showing the DEGs from the comparison of HMGA2hi to control cells **(b)**. Those DEGs with an FDR < 0.05 are depicted as red dots, and the top 10 down- and up-regulated genes are annotated. Boxplot of two stem cells markers, *HMGA2* (**c**) and *CSPG4* (**d**), that are significantly induced (FDR<0.05) in the HMGA2hi cells compared to control cells (n = 3 each). Gene expression is shown as log_2_CPM. Gene set enrichment analysis (GSEA) using the cell type signature gene sets showing two upregulated stem cell-related pathways in HMGA2hi cells compared to control cells (**e**). Gene count and significance level are shown by the size and color of each circle.

### HMGA2hi cells share a similar transcriptomic profile with HMGA2F

Disease ontology analysis using DisGeNET database (34) of HMGA2hi compared to control cells showed enrichment of multiple diseases associated with fibrosis, including renal fibrosis (Unified Medical Language System: C0151650), chronic kidney disease (C1561643), lung diseases (C0024115), and fatty liver disease (C4529962) (Figure 3a). In addition, the uterine fibroids gene set (C0042133) was also enriched by the overexpression of *HMGA2*. To determine if the transcriptomic profile of HMGA*2*hi cells is similar to uterine fibroids, we sequenced RNA from *HMGA2* fibroids (HMGA2F) and myometrium from non-fibroid patients (MyoN). The transcriptomes of the tissues separated in PC1 with 41% of variance (Figure 3b) in the PCA plot. 12,929 DEG, 6,788 downregulated and 6,141 upregulated including *HMGA2* (Log_2_FC = 11.4, FDR = 3.61 x 10^-34^), were found in HMGA2F compared to MyoN samples (Table S3). A Venn diagram of the DEGs from the HMGA2hi vs control cells and from the HMGA2F vs MyoN tissues showed a significant overlap with 61% of the DEGs (n = 434) from the cells also found in the tissue comparison HMGA2F vs. MyoN (Figure 3c).

**Figure 3.**
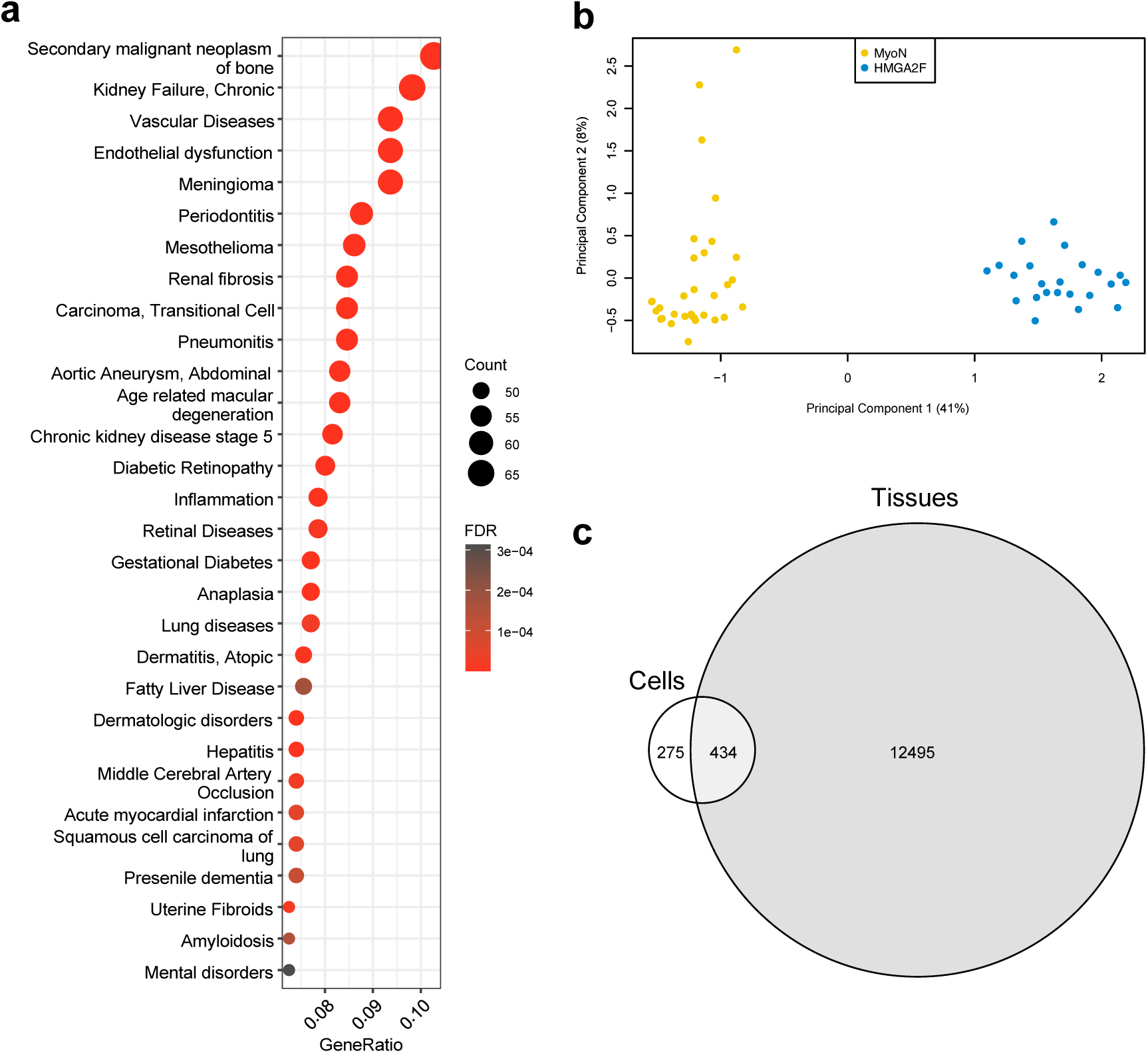
*HMGA2*hi cells have a similar transcriptomic profile with HMGA2 fibroid tissues. Disease-associated gene analysis using the DisGeNET database showing the top 30 significantly enriched pathways (FDR < 0.05) (**a**). Gene count and significance level are shown by the size and color of each circle. Principal component (PC) plot analyses for PC1 vs PC2 show samples from MyoN (n = 32) and HMGA2F tissues (n = 24) in two-dimensional space. Each dot represents one tissue sample (**b**). Venn diagram illustrate the significant (hypergeometric test; p = 2.2 x 10^-22^) overlap of DEGs between the HMGA2hi vs. control cells and the HMGA2F vs. MyoN tissue samples (**c**).

### *HMGA2*hi cells and HMGA2F tissue had dysregulated extracellular matrix

Overrepresentation analysis of gene ontology molecular functions (35) revealed co-regulated significant pathways enriched in both cells (HMGA2hi compared to control) and tissue (HMGA2F compared to MyoN). These co-regulated pathways included functions related to ECM, cell adhesion, stem cell differentiation, and muscle development (Figure 4a). To assess ECM dysregulation in both cell lines and primary cells, 3D culture followed by trichrome staining analyses were conducted. Spheroids derived from primary fibroid cells exhibited well-defined round structures, and collagen quantification revealed significantly higher deposition in fibroid spheroids compared to spheroids from MyoN cells (Figure 4b and 4c). Similarly, spheroids derived from HMGA2hi cells displayed well-defined structures and exhibited significantly elevated collagen deposition compared to their respective controls.

**Figure 4.**
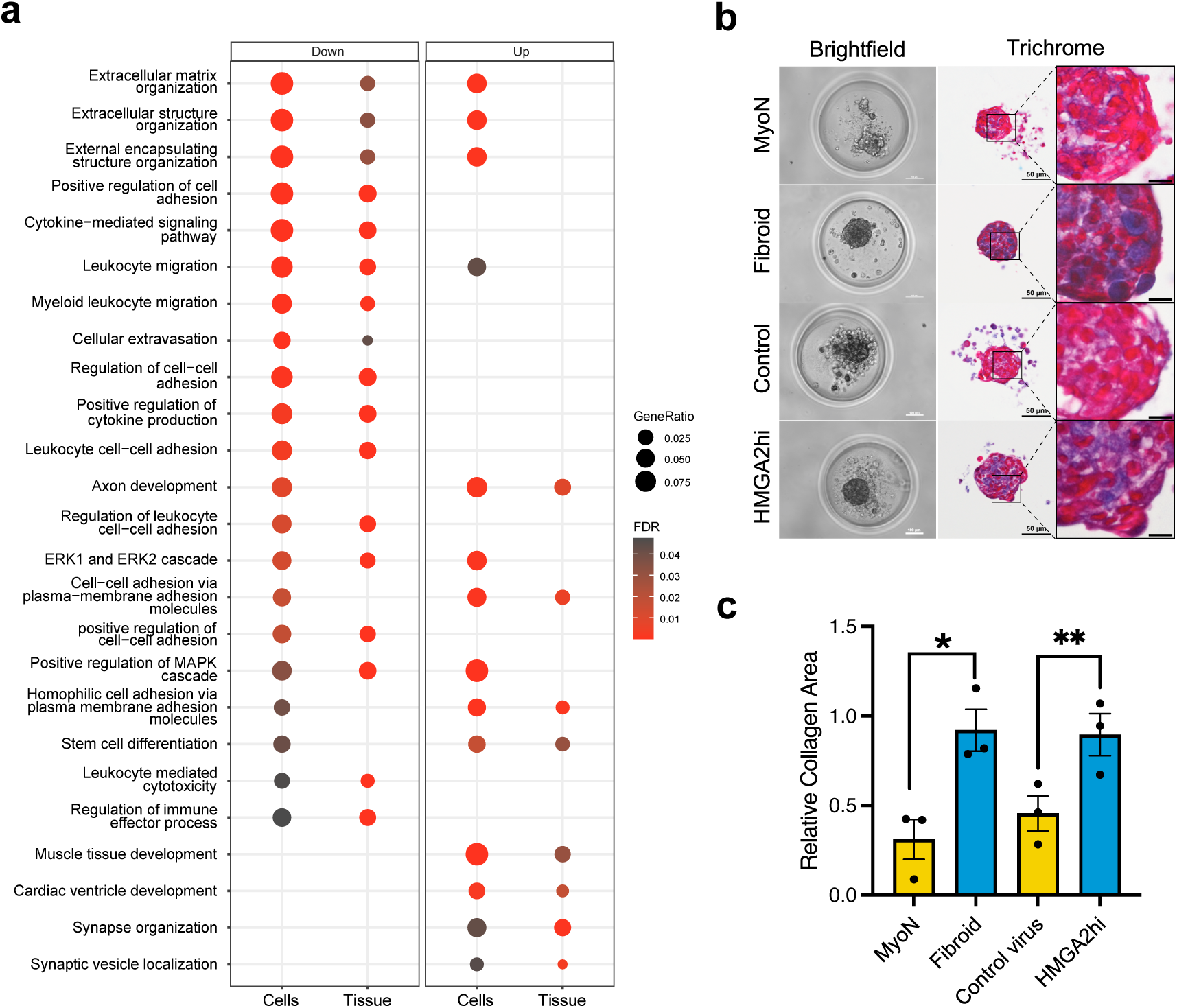
*HMGA2* overexpression in MyoN cells modulates extracellular matrix-associated pathways similarly to HMGA2F tissue. Gene ontology (GO) analysis using the biological processes (BP) gene sets showing the top 25 down- and up-regulated pathways in both cells (HMGA2hi vs. control) and tissues (HMGA2F vs. MyoN) (**a**). Gene ratio and significance level are shown by the size and color of each circle. Representative brightfield and trichrome-stained cross-section images of spheroids from primary cells and cell lines (**b**). The scale bar for the trichrome staining at higher magnitude is 10 µm. Collagen is stained in blue and cytoplasm in red. Relative collagen area, ratio of the blue/red stain, was calculated using the trichrome-stained cross sections from primary cells and cell lines (n=3 and error bars = sem) (**c**). *p<0.05, **p<0.01; by two-tailed student t test.

## Discussion

The clonal origin of fibroids from a mutated myometrial stem cell is the prevailing hypothesis for the etiology of the disease (21,22). In this study, we explored an alternative hypothesis, i.e., that fibroids could arise from a mutated differentiated myometrial cell (Figure 5). Since elevated *HMGA2* expression in adult tissues is frequently encountered in neoplastic processes, it is thought to be a key driver of tumorigenesis, but the mechanisms involved are largely unknown. HMGA2 plays a crucial role in embryonic development and stem cell maintenance (13). While its expression is generally low in most normal adult tissues (14), it is notably elevated in some tumors (13,15,16), including HMGA2 fibroids, one of the major fibroid subtypes (17,18). We exploited these HMGA2 characteristics to investigate whether its overexpression was sufficient to induce a fibroid- like phenotype in myometrial cell culture models. We found that overexpression of *HMGA2* induces a more stem cell-like phenotype in HMGA2hi cells characterized by decreased cell proliferation, enhanced colony formation, and enhanced differentiation into osteocytes and adipocytes compared to control cells, all of which support a role for HMGA2 in inducing stem cell properties in myometrial cells. Whether similar stem cell like properties can be induced in myometrial cells with other fibroid subtype genetic alterations, for example with *MED12* mutation or *HMGA1* overexpression, remains to be determined.

**Figure 5.**
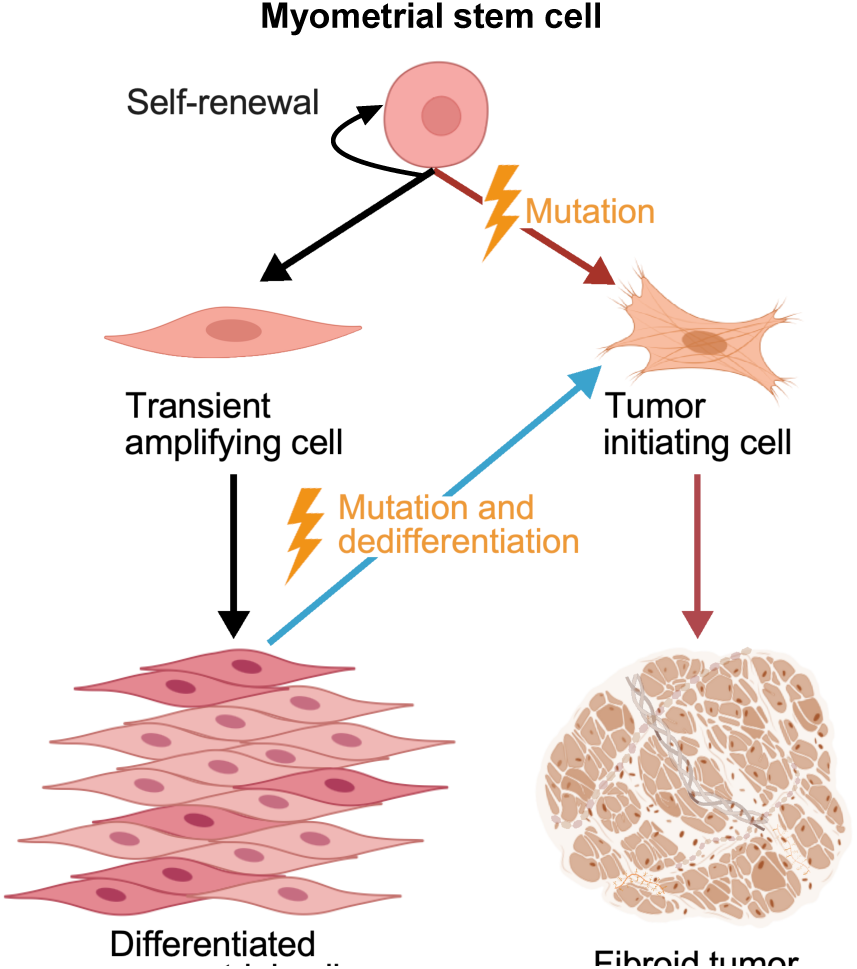
Proposed models for uterine fibroid development. Black arrows represent the traditional physiological origin of differentiated myometrial cells from a myometrial stem cell. Red arrows depict the previously proposed origin for fibroid development from a mutated myometrial stem cell. The blue arrow indicates our alternative hypothesis, suggesting fibroid tumor initiation through the dedifferentiation of myometrial cells into a more plastic tumor-initiating cell. Created with BioRender.com.

Increased colony formation in myometrial cells overexpressing *HMGA2* compared to control cells has been previously reported (12), but whether the parental cells used in the study were first depleted of putative myometrial stem cells was not described. Contradictory findings regarding cell proliferation were also noted with this previous report and the current study. While our study indicated decreased proliferation, Li, *et al.* reported a higher Ki67 proliferation index in *HMGA2*-overexpressing cells compared to the control cells. This discrepancy may be attributed to differences in the methods used to assess proliferation. In our study, we employed a continuous monitoring approach over a 10-day period in a 2D model, whereas the study by Li et al. assessed Ki67 staining at a specific time point in a 3D culture. The significance of this discrepancy is not clear and requires further investigation.

Transcriptomic analysis of HMGA2hi cells further revealed increased stemness, as evidenced by a significant increase in the expression of another mesenchymal marker, *CSPG4* (26), along with enrichment for two stem-related pathways. Several studies have demonstrated the capability of myometrial and fibroid stem/progenitor cells to differentiate into various cell types (26,32,36). However, to our knowledge, no previous reports have investigated whether myometrium and fibroid cells depleted for stem/progenitor cells retain the ability to differentiate into other cell types. Our findings suggest that the myometrial cell line can differentiate into adipocytes, and notably, can differentiate into osteocytes only when *HMGA2* is overexpressed, suggesting that *HMGA2*, a gene expressed by stem cells, enhances cell plasticity in myometrium cells.

In addition to the acquisition of the stem-like properties induced by *HMGA2* overexpression in the myometrial cell line, we have also uncovered significant transcriptomic alterations associated with *HMGA2* overexpression, indicating a pivotal role for HMGA2 in modulating key cellular processes in fibroid cells. HMGA2hi cells showed a similar transcriptomic profile to HMGA2F based on DEGs. Disease ontology analysis utilizing the sequencing data unveiled that HMGA2hi cells were enriched for the uterine fibroid gene set and over 60% of the DEGs in the HMGA2hi cells overlapped with those identified in HMGA2F tissues. This convergence in transcriptomic signatures between HMGA2hi cells and HMGA2F tissues underscores the potential role of *HMGA2* dysregulation in driving fibroid pathogenesis and further supports its involvement in modulating gene expression patterns associated with fibroid development. Mas *et al*. reported that overexpression of the truncated form of *HMGA2* in primary myometrial side population and primary myometrium cells resulted in the formation of leiomyoma-like tissue with similarities, including ECM content, to human fibroids in a xenotransplantation *in vivo* model (11). Dysregulated ECM pathways were also found in the gene ontology analysis in both HMGA2hi cells and HMGA2F tissues compared to their respective controls. Additionally, collagen deposition was significantly increased with HMGA2hi cells in a spheroid model, mimicking observations made using primary fibroid tissue cells where collagen deposition is a key histological characteristic of uterine fibroids (37).

In conclusion, *HMGA2* emerges as a central player in the pathogenesis of uterine fibroids, influencing cellular behavior and gene expression profiles. Our study sheds light on the intricate interplay between HMGA2 and fibroid development, revealing its capacity to induce a stem cell-like phenotype, alter transcriptomic profiles leading to significant changes in ECM pathways, and mirror observations in fibroid tissues, all of which underscores its role in driving fibroid pathogenesis. Further research is needed to investigate if MED12mt cells exhibit similar characteristics, confirming the theory that the mutation leads to a stem cell-like phenotype. Overall, our study highlights the significance of HMGA2 in uterine fibroid biology and lays the groundwork for future studies aimed at improving clinical management and treatment outcomes for individuals affected by this condition.

### Study approval

The use of human tissue specimens was approved by the Spectrum Health Systems and Michigan State University Institutional Review Boards (MSU IRB Study ID: STUDY00003101, SR IRB #2017-198) as secondary use of biobank materials.

### Author contributions

Experimental design (ENP, JMT), collected data and performed experiments (ENP, TJC, LP, AB, MAOB), analyzed data (ENP, JMT), wrote/reviewed manuscript (ENP, TJC, LP, AB, MAOB, ATF, JMT).

## Acknowledgments

We would like to thank the patients who consented for the study, the Spectrum Health Systems Universal Biorepository staff, and the Van Andel Research Institute Genomics Core for their help with the study. This work was supported in part through computational resources and services provided by the Institute for Cyber-Enabled Research at Michigan State University.

