## Supplementary Figure for "Unraveling the Molecular Landscape of Uterine Fibroids, Insights into *HMGA2* and Stem Cell Involvement"

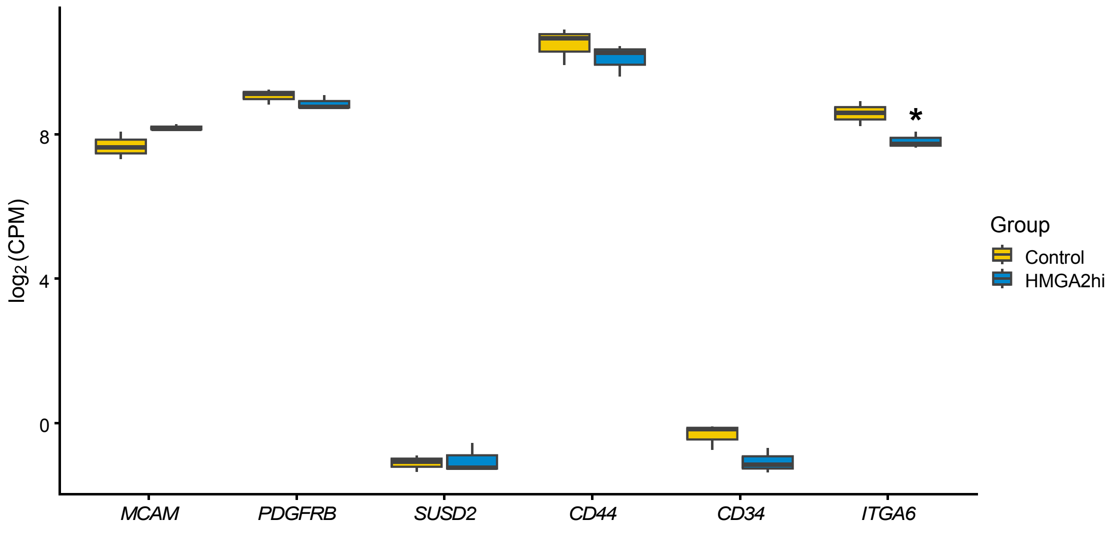


**Figure S1. Other stem cell markers in HMGA2hi and control cells.** Boxplot of stem cells markers from the RNA sequencing results in the HMGA2hi cells and to control cells (n = 3 each), *: FDR<0.05.
